# Numerical Analysis of the Immersed Boundary Method for Cell-Based Simulation

**DOI:** 10.1101/071423

**Authors:** Fergus R. Cooper, Ruth E. Baker, Alexander G. Fletcher

## Abstract

Mathematical modelling provides a useful framework within which to investigate the organization of biological tissues. With advances in experimental biology leading to increasingly detailed descriptions of cellular behaviour, models that consider cells as individual objects are becoming a common tool to study how processes at the single-cell level affect collective dynamics and determine tissue size, shape and function. However, there often remains no comprehensive account of these models, their method of solution, computational implementation or analysis of parameter scaling, hindering our ability to utilise and accurately compare different models. Here we present an effcient, open-source implementation of the immersed boundary method (IBM), tailored to simulate the dynamics of cell populations. This approach considers the dynamics of elastic membranes, representing cell boundaries, immersed in a viscous Newtonian fluid. The IBM enables complex and emergent cell shape dynamics, spatially heterogeneous cell properties and precise control of growth mechanisms. We solve the model numerically using an established algorithm, based on the fast Fourier transform, providing full details of all technical aspects of our implementation. The implementation is undertaken within Chaste, an open-source C++ library that allows one to easily change constitutive assumptions. Our implementation scales linearly with time step, and subquadratically with mesh spacing and immersed boundary node spacing. We identify the relationship between the immersed boundary node spacing and fluid mesh spacing required to ensure fluid volume conservation within immersed boundaries, and the scaling of cell membrane stiffness and cell-cell interaction strength required when refining the immersed boundary discretization. This study provides a recipe allowing consistent parametrisation of IBM models.

## 1. Introduction

The collective dynamics of populations of cells play a key role in tissue development and self-renewal, as well as in disease. Mathematical modeling of these systems is challenging due to the wide range of behaviours displayed over different time and length scales, necessitating increasingly sophisticated ‘multiscale’ approaches [4]. Such models seek to gain insight into emergent behaviours where the coordinated action of cell-scale processes, such as the localisation of membrane-bound planar cell polarity proteins, can combine to effect striking tissue-level morphogenetic changes, such as convergent extension, in a variety of developing epithelial tissues [31].

Molecular and live-imaging techniques allow tissues to be probed at ever finer scales, supporting the use of modelling frameworks within which hypotheses spanning from the subcellular to the tissue scales may be tested. A range of such models have recently been developed, from vertex models that approximate each cell geometrically by a polygon or polyhedron representing the cell’s membrane [8] to subcellular element models that allow for more arbitrary cell shapes [25, 26]. Yet a firm mathematical foundation for the analysis of such models, which is required for confidence in the conclusions drawn from them, remains lacking. To help address this, we present a detailed examination of the immersed boundary method (IBM), which forms the basis for one such class of model, and a computational implementation thereof, designed to study interacting populations of eukaryotic cells.

The IBM is a numerical method for simulating the dynamics of one or more elastic membranes immersed in a viscous Newtonian fluid. It was first developed by Peskin to investigate flow patterns around heart valves [16]. The model is formed from two coupled components: elastic boundaries representing, for instance, heart valves or cell membranes, and a fluid extending over the entire spatial domain. The elastic boundaries exert a force on the fluid, which induces a flow that, in turn, causes the boundaries to move. In the context of interacting cell populations, each immersed boundary may be thought of as representing the membrane of an individual cell or, more generally, structures on smaller or larger scales such as intracellular detail [6] or multicellular regions of tissue [5]. Inter- and intra-cellular interaction terms, which represent phenomena such as cortical tension in the cell membrane and the action of adhesive transmembrane proteins, are specified as explicit forces acting between discrete locations on each immersed boundary. A schematic of parts of three neighbouring immersed boundaries is shown in Figure 1. The set of such interactions defines, at any given time, a resultant force acting on the membranes. This resultant force is applied to the fluid, which induces a flow. This flow carries the membranes along with it, thereby updating the positions of the boundaries. Thus, the role of the fluid is to provide a mechanism by which the boundary locations are updated; a more detailed discussion of this mechanism is presented in Section 2.

**Fig. 1.**
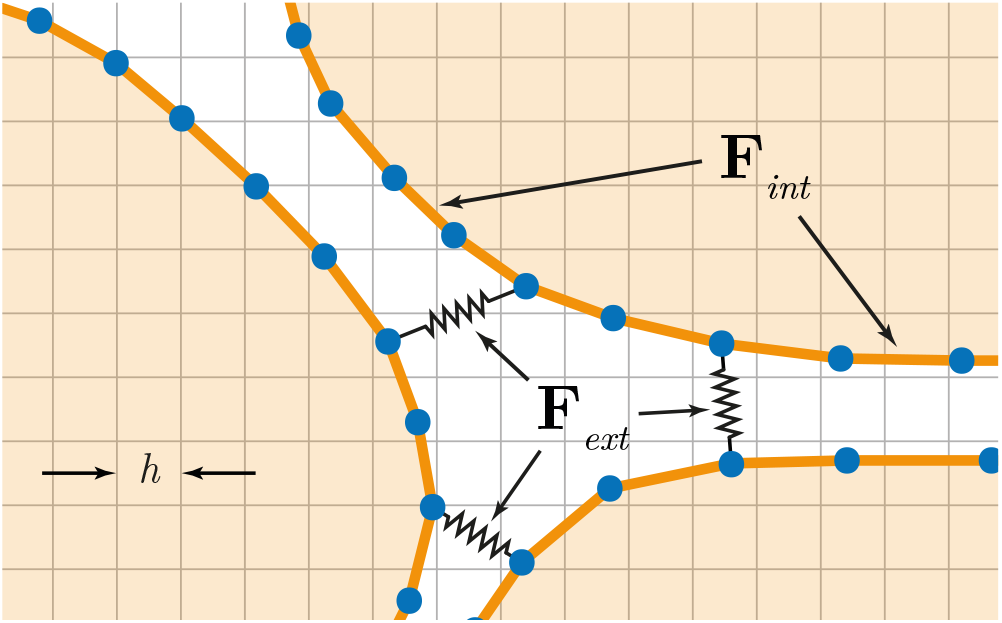
Schematic of immersed boundaries. Circular nodes represent an off-lattice discretization of the immersed boundary contours. The regular grid behind the boundaries represents points on which a viscous Newtonian fluid, ubiquitous across the domain, is discretized. Adhesion links, specified as explicit force terms, exist between nodes within each immersed boundary, as well as between neighbouring boundaries. The terms h, **F**_int_, and **F**_ext_ are defined in the main text.

The IBM has several features that make it well suited to modelling the collective dynamics of cell populations. First, and most importantly, the shape of cell boundaries can be represented with arbitrary precision. This enables investigation of processes at a subcellular scale, while allowing cell shapes to be an emergent property rather than a constraint of the model, in contrast to other approaches such as vertex models [20, 27] and spheroid models [10]. Second, volume is preserved within any given closed contour of the fluid, unless specifically altered by fluid sources or sinks. Thus, the IBM allows for the study of regulated processes that affect cell size, such as cell growth, shrinkage, division and death. Third, implementations of the IBM typically have a small number of parameters. As shown in Section 3, the fluid dynamics depend only on the Reynolds number, while cell mechanical interactions are usually modelled by means of simple forces, such as linear springs. This opens the possibility of calibration against biological data; Rejniak, for instance, has demonstrated this by successfully parametrising an IBM implementation with numerical values estimated from various experimental studies [23]. Finally, unlike numerical schemes that employ structured or unstructured grids conforming to the immersed body, in the IBM the fluid is discretized using a regular Cartesian grid that may be generated with ease. This allows a relatively simple numerical scheme, discussed in Subsection 4.8, which has a fairly straightforward and efficient computational implementation, and enables the use of a fast and direct spectral method for computing the fluid flow.

Several previous studies have detailed aspects of the IBM, including a thorough treatment of the underlying mathematics by Peskin [17]. Biological applications include those by Rejniak et al., who use an IBM implementation to investigate the growth of solid tumours under differing geometric configurations, initial conditions, and tumour progression models [22, 23]. The same authors have investigated the mechanics of the bilayer of trophoblasts in the developing placenta [24]. Dillon and Othmer use an IBM to model spatial patterning of the vertebrate limb bud [5], and an IBM framework for tackling general morphogenetic problems is presented by Tanaka et al. [29]. Cell deformation is investigated by several authors; by Jadhav and colleagues in the context of cell rolling [11] and by Bottino in the context of passive actin cytoskeletal networks [1]. A review by Mittal and Iaccarino gives excellent background on the method and cites many other examples of its use across various application areas [14].

While, collectively, these papers provide an excellent overview of the IBM and several implementations thereof, there remains no comprehensive account of the model, method of solution, computational implementation or analysis of parameter scaling. The aim of this work is therefore to provide comprehensive details of an IBM implementation aimed specifically at describing the collective dynamics of multicellular tissues. We provide a free, open-source implementation of the IBM complete with example simulations: we build on the established Chaste library [13, 19] to ensure that the code is robust and well-tested; we present the code necessary to reproduce all figures in this paper; and we conduct a thorough numerical analysis detailing how parameters scale with respect to each other in order to build a recipe allowing consistent parametrisation of models. The remainder of this paper is structured as follows. Sections 2 to 4 give details of the IBM, its discretization, and a numerical solution using a fast Fourier transform algorithm. Section 5 outlines the C++ implementation in Chaste. Section 6 details a numerical analysis demonstrating that the computational implementation converges, and elaborating on how parameters scale relative to each other. Section 7 concludes with a discussion of the choices made in our implementation, and future work in this area.

## 2. Immersed boundary method formalism

Consider a viscous Newtonian fluid, with velocity **u** = **u**(**x**) = **u**(*x, y*),flowing in a two-dimensional, doubly periodic domain Ω = [0, *L*] × [0, *L*]. The fluid motion obeys the Navier-Stokes equations

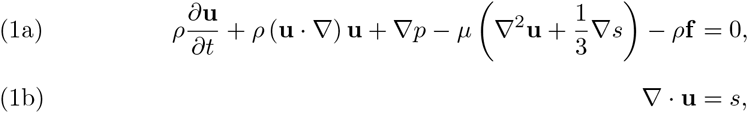

where *ρ* and *µ* are the fluid density and viscosity, respectively, and are both assumed constant; *p* is the pressure field; **f** is the force per unit area acting on the fluid; and *s* is the fluid source field, representing the proportional volume change per unit time. The periodic boundary conditions enforce **u**(*x*, 0) = **u**(*x*, *L*) and **u**(0, *y*) = **u**(*L*, *y*), for 0 ≤ *x*, *y* ≤ *L*.

We next consider a set of *N* non-overlapping closed curves in the fluid, which we will refer to as immersed boundaries, and which we think of as representing cell membranes. We associate upper-case Roman indices with the immersed boundaries, and lower-case Roman indices with the fluid domain Ω. Let 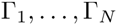 denote the collection of immersed boundaries, and let each immersed boundary Γ_*k*_ be parametrised by γ_*k*_. Further, let

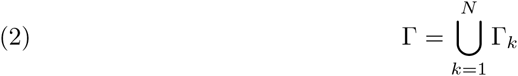

denote the union of these immersed boundaries, parametrised by γ, which is composed of 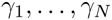 in the natural way. Let **X** = **X**(γ_*k*_, *t*) denote the coordinates of the k^th^ immersed boundary and let **X** = **X**(γ, *t*) be the combined coordinates of all immersed boundaries.

We denote the resultant force acting on the immersed boundaries by **F** = **F**(γ, *t*). The precise functional form of the resultant force **F** varies with application, and is formulated in Section 4. We relate the resultant force on the immersed boundaries to the body force acting on the fluid through the relation

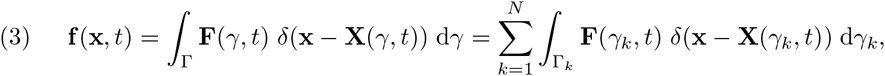

where δ(·) denotes the Dirac delta function. The force on the fluid at location **x** thus vanishes away from the immersed boundaries, and equals the resultant force **F** at location **X** on an immersed boundary precisely at **x** = **X**.

The immersed boundaries are assumed to move due to the fluid flow without slipping, so that a point along Γ moves at precisely the local fluid velocity:

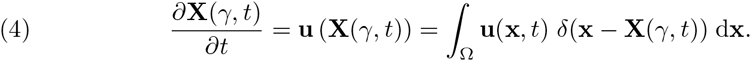

Thus, the velocity of an arbitrary immersed boundary point **X**(γ) is equal to the velocity of the fluid at **x** = **X**.

The source field, *s*, is considered to be a finite linear combination of individual point sources. The number, location, and strength of each source is formulated in Section 4, but for now we consider s as an arbitrary (but known) scalar field.

## 3. Non-dimensionalization

We non-dimensionalize the model to reduce the number of parameters and allow us to estimate the relative importance of each term. For the Navier-Stokes equations, we introduce the standard choices for viscous dynamics: a length scale, *L*; velocity scale, *U*; time scale, *L*/*U*; pressure scale, U*µ*/*L*; source scale, *U*/*L*; and force scale, *U*^2^/*L*. Substituting the rescaled variables and operator

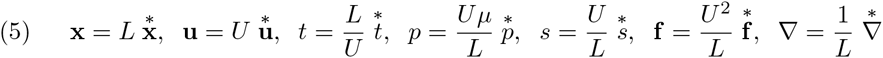

into Equations (1a) and (1b) and dropping the stars yields

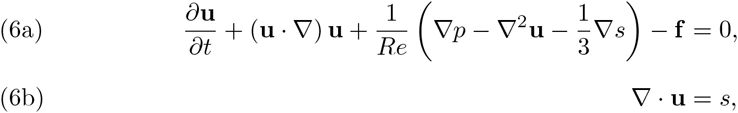

where *Re* = *ρLU*/*µ*, the Reynolds number, represents the ratio of inertial to viscous forces. At very low Reynolds number it is appropriate to take the limit *Re* → 0 in Equation (6a) and, assuming the body force, **f**, to be of order 1/*Re*, obtain the Stokes equations

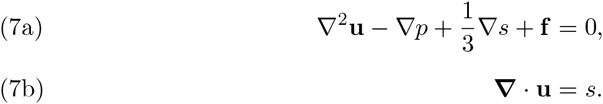

Note that we assume 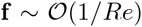 since otherwise no flow would be induced by the force on the immersed boundaries.

Small scale systems typically exhibit low velocities, and thus Reynolds numbers for biological regimes can be very small. Small swimming organisms, for instance, may experience Reynolds numbers as low as 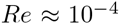 [21]. Tanaka et al. [28] estimate Reynolds numbers for the fluid-like properties of embryonic tissues as 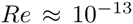 using assumptions of 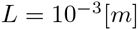, 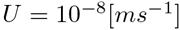 and 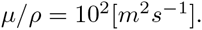. Rejniak et al. [24] arrive at 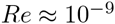 by considering the length scale to be the size of a large cytotrophoblastic cell (20*µ*m) and a characteristic velocity of 30*µ*m in 24 hours.

Equations (7a) and (7b) are computationally less expensive than the full Navier-Stokes to solve; for example, their linearity permits the use of efficient Green’s function methods [3]. This raises the question of the circumstances under which it is appropriate to assume Stokes flow for the IBM fluid component, as described in [2, 12]. Here, we choose to solve the full Navier-Stokes equations, the reasons for which are discussed in Section 7, while keeping in mind that there are particular simulations for which the reduced problem may be suitable and computationally less expensive to solve.

Having chosen to solve the non-dimensional Navier-Stokes equations (6a) and (6b), we non-dimensionalize (Equations 3) and (4) using the rescaled parameters

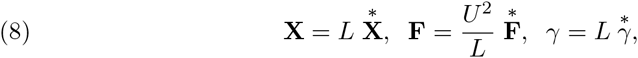

dropping the stars, as before.

## 4. Discretization

We solve the coupled problem, consisting of (Equations 3), (4), (6a), and (6b), numerically, as follows. The immersed boundaries are discretized into a finite union of points (small circles in Figure 1) that we call nodes. The fluid domain is discretized onto a regular Cartesian grid (square lattice in Figure 1) that we refer to as the mesh. In our non-dimensional coordinates, 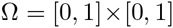 is discretized with *N* × *N* grid points with mesh spacing *h*. We must also discretize Equation (3) relating the force **F** on the immersed boundaries with the body force **f** acting on the fluid, and Equation (4) relating the fluid and node velocities.

In the following, time is discretized in steps of Δ*t*, and we refer to an arbitrary function 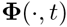 at the *n*^th^ time step by 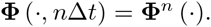.

### 4.1. Discrete Dirac delta function

In the discretized system, the fluid and immersed boundaries interact only via a discrete version of the Dirac delta function. To approximate the Dirac delta function on the discrete mesh, we require a function with finite support for which, when interpolating between the immersed boundary and fluid domains, the contributions at each fluid mesh point in the support sum to unity. Various such functions have been proposed, of which four examples from different IBM implementations are detailed by Mittal and Iaccarino [14].

Here, we make the common choice of a trigonometric function, used in several other IBM implementations [5, 23, 24], given by

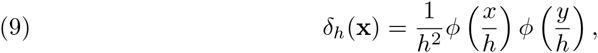

where *h* is the mesh spacing, and the function *ϕ* is given by

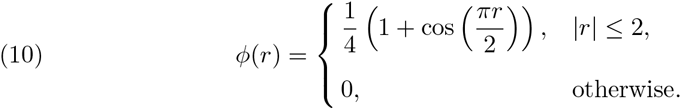

This choice of *ϕ* differs from, but takes extremely similar numerical values to, that derived and used by Peskin [17]. Given the numerical similarity, the choice of function is unlikely to make much practical difference, and we have found the version presented here to be less computationally expensive to compute (see Section 7).

We also note that, due to the bounded support of both functions, *δ*_*h*_(**x**) will only ever be non-zero at the 4×4 mesh points closest to any given node. The choice of support size is discussed by Peskin [17], and is made purely on computational grounds: one could choose a delta function approximation with wider support, but each node on an immersed boundary would then interact with many more mesh points, slowing down the computation.

### 4.2. Discretization of immersed boundaries

We discretize each immersed boundary Γ_*k*_ by a set of *N*_*k*_ nodes whose positions are given by 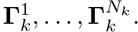. We suppose that these nodes are initially spaced equally along the original parameter range 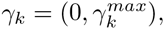, so that the length element 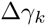 associated with the *k^th^* immersed boundary is equal to the initial node spacing, 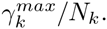. Since we impose the condition that each immersed boundary forms a closed contour we have 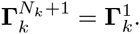

### 4.3. Discrete force relations

We are now in a position to define the resultant force **F** acting on the immersed boundaries. The discretization treats **F** as the union of a finite set of point forces given by the resultant force on each node in each immersed boundary.

We will consider the resultant force on each node as the combination of two types of force: `internal’ forces, which depend on the positions of other nodes in the same immersed boundary; and `external’ forces, which depend on the positions of nodes in different immersed boundaries. Here, we introduce specific choices for the force terms to represent both the mechanical properties of the actomyosin cortex of a cell and the protein-protein interactions between neighbouring cells. We represent both these mechanical interactions by linear springs, following previous IBM implementations [5, 22, 23, 24, 29], although different functional forms could, in principle, be chosen.

Internal forces represent the contractile properties of a eukaryotic cell’s actomyosin cortex, which we describe by connecting each node to its neighbours by a linear spring of stiffness *κ*_*int*_ and rest length *l*_*int*_. The internal force acting on node 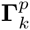 is thus given by

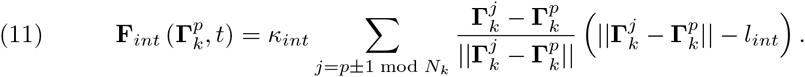

External forces represent the adhesive properties of transmembrane proteins, such as integrins and cadherins, linking neighbouring cells. We assume that any node in an immersed boundary is connected to all nodes in different immersed boundaries that are situated within a distance *d*_*ext*_ by a linear spring with stiffness *κ*_*ext*_ and rest length *l*_*ext*_. The external force acting on the node 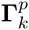 is given by

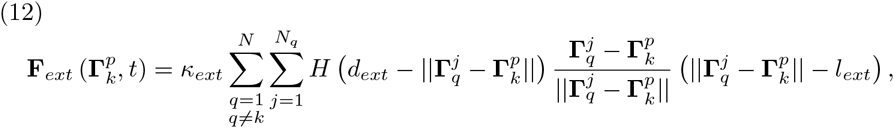

where the outer sum runs over all other immersed boundaries, the inner sum runs over the *N*_*q*_ nodes in boundary *q*, and *H*(·) is the Heaviside step function. Our choice of linear spring interactions is motivated primarily by their ease of implementation and low computational overhead (see Section 7), although in our software implementation the user is free to define their own functional forms.

The total force **F** on a node is given by the sum of the internal and external forces,

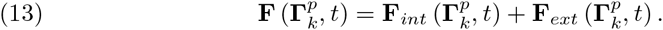

### 4.4. Discretization of the Navier-Stokes equations

Due to the periodicity of the spatial domain, we employ a fast Fourier transform algorithm to solve Equations (6a) and (6b) numerically. We use the following numerical scheme, described first by Peskin and McQueen [18] and later, with the addition of fluid sources, by Dillon and Othmer [5], where the sums are taken over the two dimensions, *d* ϵ {1,2}:

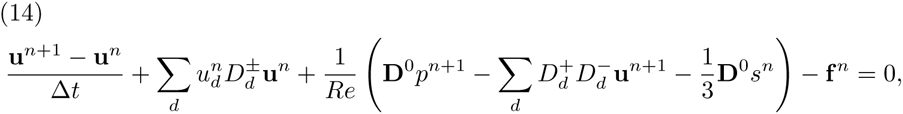

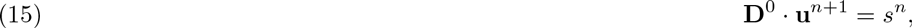

where the forward and backward divided difference operators, 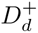and 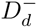, the vector of central divided difference operators, **D**^0^, and the upwind divided difference operator, 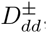, are defined by

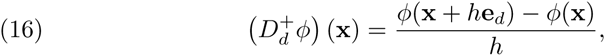

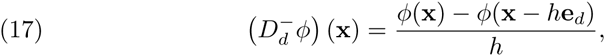

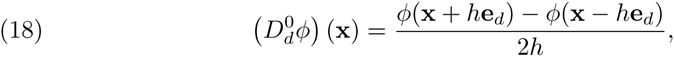

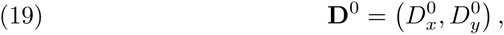

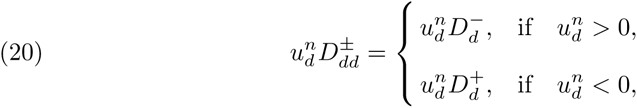

respectively. Here, **e**_*d*_ denotes the unit vector in the d^th^ dimension.

### 4.5. Discretization of force relation

We discretize Equation (3), relating the force on the fluid to the force on the immersed boundaries, as follows. For each point **x** in the fluid mesh, we sum the force contributions from every immersed boundary node using the discrete delta function to assign the appropriate weight,

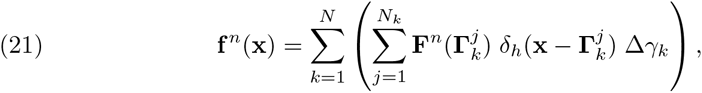

where the outer sum runs over the *N* immersed boundaries, the inner sum runs over the *N*_*k*_ nodes in the *k*^th^ immersed boundary, and Δγ_*k*_ is the length element associated with the *k*^th^ immersed boundary.

### 4.6. Discretization of position-updating relation

For simplicity, we discretize Equation (4) using a forward Euler scheme. At the *n*^th^ time step, a given immersed boundary node 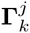 is displaced by 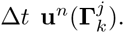. Since, in general, 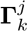 will not coincide with a fluid mesh point, the value 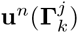 is an interpolation of the 4×4 fluid mesh points closest to 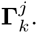. The discretized relation for updating node locations is therefore given by

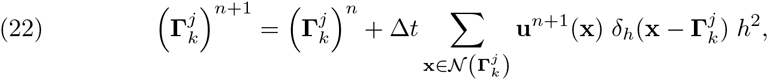

where 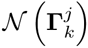 represents the 4 × 4 fluid mesh points nearest 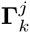 (the only points with non-zero contributions, due to the implementation of *δ*_*h*_).

### 4.7. Discretization of fluid sources

The source term s allows individual regions enclosed by contours in the fluid domain to increase or decrease in volume. In the absence of *s*, due to the volume conservation property of the IBM, the quantity of fluid within any given closed contour remains fixed. In the context of simulating multicellular biological systems, the source term *s* allows the modulation of cell size.

To allow the fluid volume within each immersed boundary to be modulated, we decompose *s* into a finite number of point sources and initially put a single source at the centroid of each immersed boundary. To ensure a constant total volume of fluid in the domain Ω, we additionally include a number of sinks (sources with a negative strength) located outside all immersed boundaries which balance the magnitude of the *N* sources associated with the boundaries.

Suppose there are *M* combined sources and sinks, 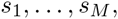, with *M*>*N*, located at the positions 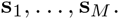. Each source *s*_*k*_ has specified strength *T*_*k*_, where PM 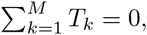, and the source field *s*(**x**) at an arbitrary fluid mesh point **x** then satisfies

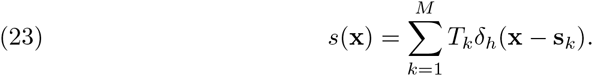

A convenient method to ensure that fluid sources always remain inside (or outside) immersed boundaries entails updating their locations in the same way as for the immersed boundary nodes,

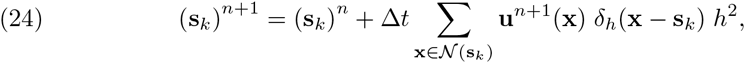

where 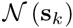 represents the 4×4 fluid mesh points nearest **s**_*k*_.

The regulation of source strengths depends on the application and on the biological process underlying the cell size change, and may, for example, be linked to a description of cell cycle progression and growth. Some examples of biological processes and their feedback on source strengths are discussed in Subsection 5.2. Note also that the number of extra `balancing sources’ is not fixed, and this is discussed in Section 7.

### 4.8. Numerical solution

We are now in a position to solve (Equations 6a) and (6b) numerically. Equation (21) allows the direct computation of **f**^*n*^, but Equation (22) requires **u**^*n*+1^, which we must compute, given **f**^*n*^, from (Equations 14) and (15).

Rearranging Equation (14) to separate the terms evaluated at different time steps yields

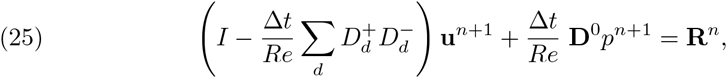

where

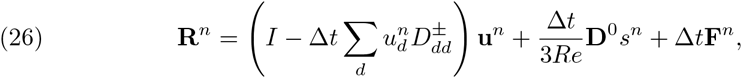

and *I* is the 2×2 identity matrix.

We solve (Equations 15) and (25) directly for **u**^*n*+1^ by applying a discrete Fourier transform (DFT) to eliminate *p*^*n*+1^. For our domain 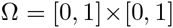 discretized using an *N*×*N* square mesh of spacing *h*, we define the DFT from the spatial coordinates (·)_*x*,*y*_ to the spectral coordinates 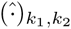 by

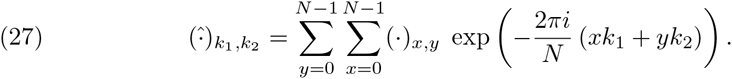

Under this transformation, (Equations 15) and (25) become

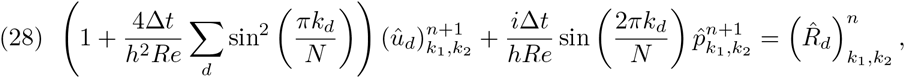

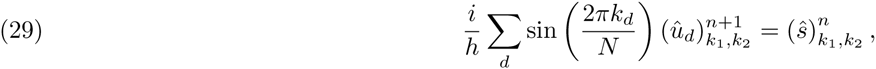

where *i* is the imaginary unit, sums are taken over dimension, 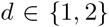, and each equation is now of a single variable so holds for *d* = 1, 2.

We substitute Equation (29) into Equation (28) to solve directly for *p*: multiplying Equation (28) by (*i*/*h*)sin(2π*k*_*d*_/*N*), summing it over the two dimensions, and rearranging for 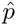 gives

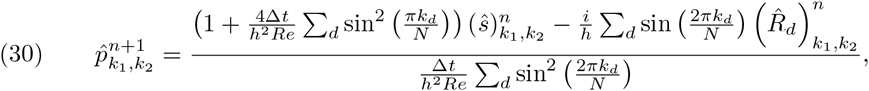

where every term on the right-hand side depends only on information available at the current time step. We can therefore substitute Equation (30) back into (Equation 28) to solve for 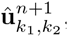, obtaining

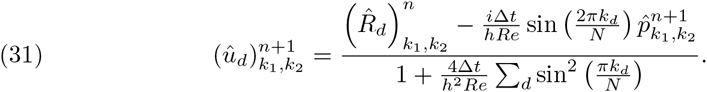

Care must be taken at the mesh points (*k*_1_, *k*_2_) = (0, 0), (0, *N*/2), (*N*/2, 0) and (*N*/2, *N*/2), where the denominator of the right-hand side of Equation (30) vanishes. At these points, however, the sine term multiplying 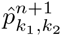 in (Equation 28) also vanishes, and we may thus solve directly for 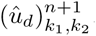. We, therefore, avoid this problem by setting 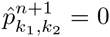 in Equation (31). Finally, having computed 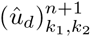, we apply the inverse DFT to obtain **u**^*n*+1^,

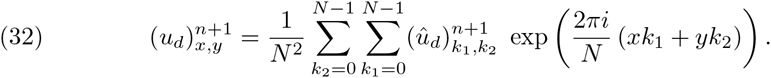

## 5. Computational implementation

In this section, we describe the time-stepping algorithm for solving the IBM model, and how it fits into the computational modelling framework Chaste. We go on to highlight some of the computational challenges addressed during the implementation of this method, and we present some benchmarking and profiling results. Finally, we detail how rule-based processes such as cell division, needed for simulating populations of cells, are implemented within this IBM implementation.

### 5.1. Chaste

We have implemented our IBM model as part of the Chaste C++ library [13, 19]. The IBM code is released as a feature branch of the latest development version of Chaste^1^, which is open source and available under the 3-clause BSD licence. Chaste is developed with a test-driven approach using the unit testing framework CxxTest^2^. Using this framework, unit tests verify the correctness of every individual method within the implementation, and simulations are themselves written as test suites. Details of how to obtain our IBM implementation, as well as code to recreate all simulations from this paper, can be found in Appendix A.

As it is written in C++, Chaste is fast and able to utilise object orientation and class inheritance, enabling modularity and easy extensibility of the code base. This structure enables the IBM to integrate with Chaste as a new example of the pre-existing class of `off-lattice simulations’, within which much of the core functionality such as simulation set up, time stepping, and cell cycle models are already implemented and thoroughly tested. In addition, new specialised functionality is built upon existing abstract classes, meaning a consistent and familiar interface is presented to existing code users.

Using the numerical method described in Section 4, we solve the IBM by iterating through the following fixed time-stepping algorithm:

1. update the cell population to take account of cellular processes including cell death, division, growth, shrinkage, and procession through the cell cycle, discussed shortly;
2. calculate the internal and external forces acting on each node, using Equation (13);
3. loop over each immersed boundary node and propagate the associated force to the fluid mesh domain, as described by (Equation 21);
4. loop ever each fluid source and propagate the associated strength to the fluid mesh domain, as described by (Equation 23);
5. solve the Navier-Stokes (Equations 6a) and (6b) using the fast-Fourier transform algorithm detailed in Subsection 4.8 to generate new fluid velocities;
6. use the new fluid velocities to update immersed boundary node and fluid source locations as described by (Equations 22) and (24).

An example of a simple simulation performed using this implementation within Chaste is visualised in Figure 2, where an elliptical immersed boundary relaxes over time towards a circular shape. The fluid flow is shown as a vector field of arrows.

**Fig. 2.**
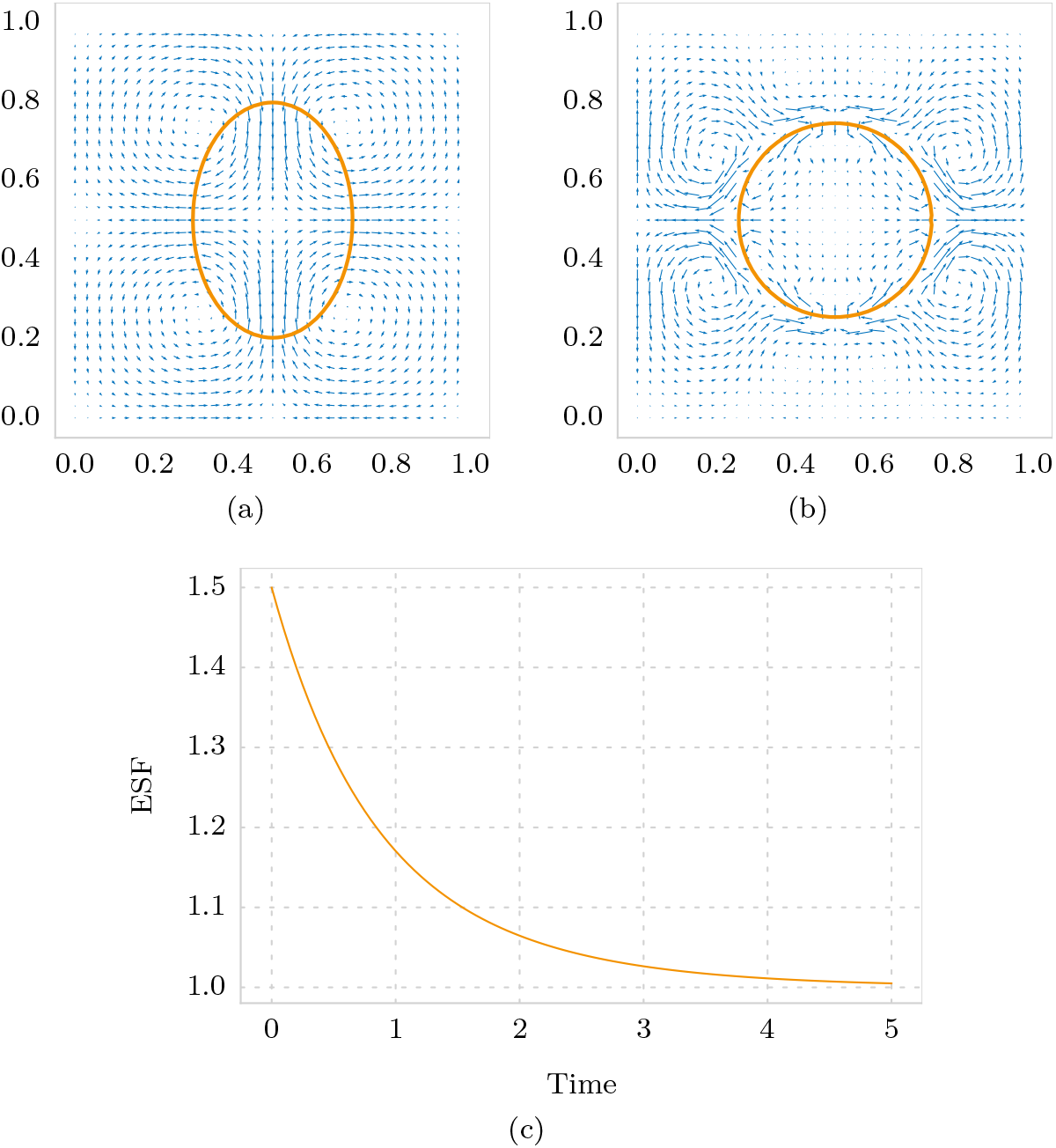
An example IBM simulation. An elliptical immersed boundary relaxing over time under the action of internal forces (Subsection 4.3), and no fluid sources. In this simulation, h = 1/32, Δt = 0:05 and N = 128 nodes. Full details of all parameters can be found in the simulation code, available as part of the test suite `TestNumericsPaperSimulations’ (see Appendix A). (a) State of the simulation after one time step, where flow is acting to reduce the elliptical immersed boundary in height and expand it in width. (b) State of the simulation after 100 time steps, where flow vanishes at the boundary. (c) Dynamics of the aspect ratio of the ellipse, quantified by its elongation shape factor (ESF; see Section 6 for details), over time.

### 5.2. Implementation of cellular processes

Sections 2 to 4 detail our IBM model and a numerical solution thereof, and together these constitute a method of solving fluid-structure interactions. In addition to this, we need the facility to include various cellular processes that occur when modelling a multicellular tissue. Such processes include regulated cell growth, division and death, and can be thought of as a collection of rules by which the properties of the immersed boundaries or fluid sources are altered, but which do not directly alter the underlying fluid problem.

An example of rule-based modification of immersed boundaries is cell division. Within Chaste, we make use of existing functionality for encoding cell cycle progression. In this framework, a cell may at some time step be deemed `ready to divide’, at which time the following scheme is employed to replace the single immersed boundary (representing the cell about to divide) with two immersed boundaries (representing the daughter cells). First, a division axis through the centroid of the immersed boundary is selected, by means of some rule chosen by the user. This rule may, for instance, select the shortest axis of the immersed boundary, or a random axis, depending on the biological assumptions particular to the scenario being modelled. Second, with the division axis fixed, the boundary is divided in two: nodes on each side of the axis define the shape of each daughter cell, with the daughter cells separated by a pre-determined fixed distance. We make the choice that each daughter cell is represented by the same number of nodes as the parent cell, and a re-meshing process instantaneously spaces these nodes evenly around the outline of each daughter cell.

We remark that this scheme defines a rule-based implementation of cell division as a process occurring during a single time step. Depending on the time scale over which the tissue is modelled, one may wish to explicitly represent pinching during cytokinesis, as implemented by Rejniak and colleagues [22, 24]. This can be achieved within Chaste, using existing functionality that allows feedback between the cell cycle and arbitrary cell properties such as a `target’ surface area that cells seek to attain. In this manner, when a cell is selected to divide, processes such as an increase in size followed by the formation of a contractile furrow could be specified (for instance, via a feedback with fluid source strengths); however, we stress that our implementation is left flexible and extendible. The modular and hierarchical nature of Chaste allows the user to easily specify appropriate cell cycles, division rules and cell property modifiers for a given biological scenario.

### 5.3. Computational effciency and profiling

The two most computationally demanding steps in our IBM implementation are solving the Navier-Stokes equations, and calculating the forces acting on the immersed boundary nodes. The former is demanding due to the calculations necessary in the finite difference scheme, the five two-dimensional DFTs per time step, and the term-by-term calculation of the pressure field. The latter is costly due to the potentially large number of pairwise interactions between nodes on neighbouring immersed boundaries that must be kept track of.

To reduce the time spent solving the Navier-Stokes equations, we ensure that all arrays storing values needed during the computation are instantiated during simulation set up, and remain in place throughout the simulation. For *N* × *N* fluid mesh points, this means permanently storing 12*N*^2^ double-precision numbers. The result of this is a drastic speed-up compared to dynamically allocating memory, with the drawback of a large memory footprint. In practical terms, this scheme puts an upper bound of *N* ≈ 4096 when running a single simulation on a desktop computer, which is not prohibitive.

To optimise the second problem of efficiently calculating pairwise interactions between nearby immersed boundary nodes, we employ a spatial decomposition algorithm [9]. The domain is broken into squares each the size of the interaction distance *d*_*ext*_, and at each time step the nodes are placed into their corresponding square. For a given node, the only possible set of interactions are then between nodes in the same or neighbouring squares. Thus, we dramatically reduce the computation necessary when *d*_*ext*_ ≪ 1.

Table 1 shows various profiling statistics for a prototype simulation of 20 cells initially arranged in an hexagonal packing. The columns of Table 1 each represent a successive doubling of the resolution of both the fluid mesh and the immersed boundary nodes. Figure 3 shows the configuration of the immersed boundaries at the start and end of the simulation corresponding to the first column in Table 1. As can be seen, solution of the fluid problem scales less well than calculating the forces; however, neither component indiviudally dominates the simulation runtime.

**Fig. 3.**
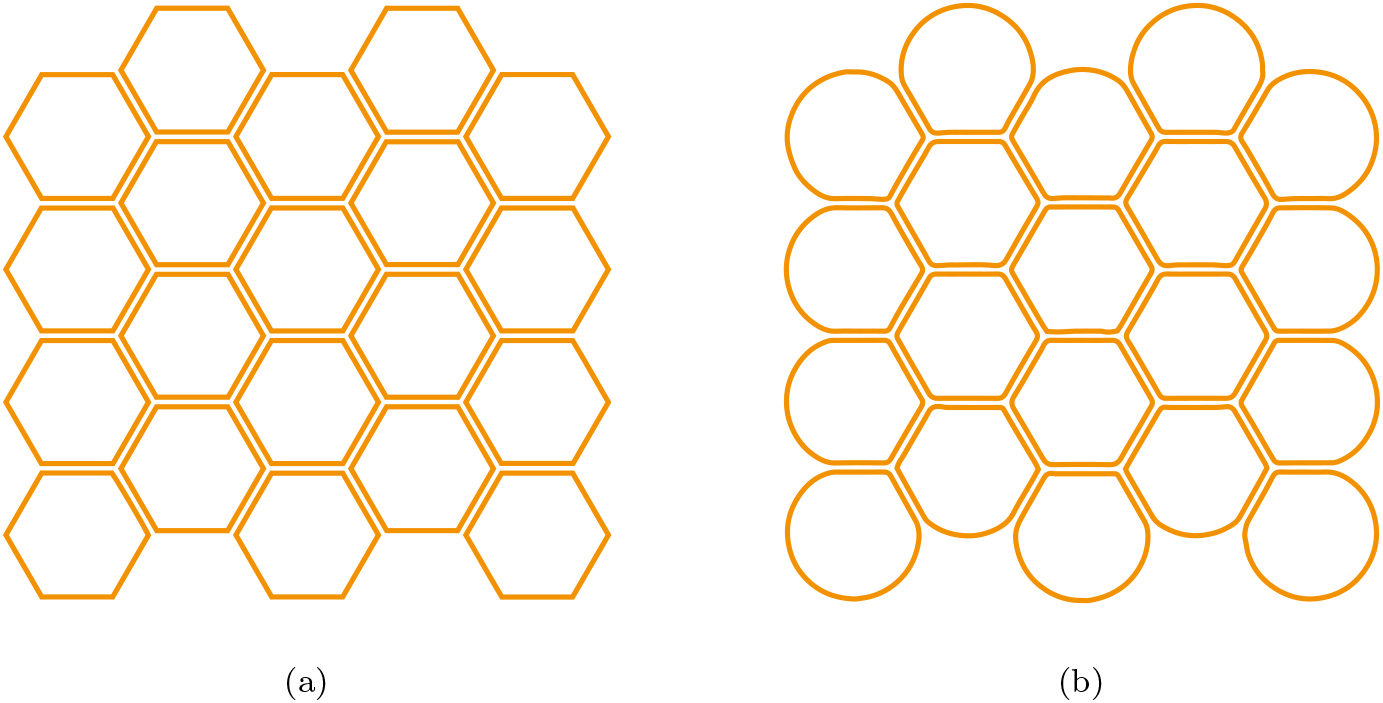
Profiling simulation. (a) The initial configuration of the immersed boundaries. (b) The configuration of the immersed boundaries after advancing 2000 time steps.

**Table 1.** Code profiling. The memory footprint, time to complete 2000 time steps, and the proportion of time spent solving the Navier-Stokes problem is presented for each of three increasingly fine simulation representations. Each simulation comprises a regular hexagonal lattice of 20 immersed boundaries, allowed to relax for the fixed number of time steps. Each boundary has 300, 600, and 1200 nodes in separate simulations with 512, 1024, and 2048 fluid mesh points, respectively. Profiling was performed on a desktop machine with an Intel Xeon E5-1650 v3 CPU and 16GiB RAM, using the GNU gprof profiler. For details of how to obtain the code for these profiling simulations, see Appendix A.

| | $h = 1/512$ | $h = 1/1024$ | $h = 1/2048$ |
| --- | --- | --- | --- |
| Approximate memory footprint (MiB) | 39 | 102 | 355 |
| Time to advance 2000 time steps (s) | 73.9 | 211 | 817 |
| Time solving the fluid problem (%) | 40.7 | 62.7 | 77.3 |

## 6. Numerical results

In this section, we run a number of simulations to demonstrate various properties of our IBM implementation. We first highlight an important relationship between the immersed boundary node spacing, Δγ_*k*_, and the fluid mesh spacing, *h*. We go on to explore how certain parameters in the IBM scale with each other, and use this to work towards a recipe by which a model of a particular biological process may be simulated. Finally, we demonstrate that the implementation converges in time step, in fluid mesh spacing, and in immersed boundary node spacing. We employ a summary statistic for an individual cell in a simulation, referred to as the elongation shape factor (ESF). For a polygon this is a dimensionless positive real number that defines a measure similar to aspect ratio. Formally, it is defined as 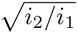, where *i*_1_ < *i*_2_ are the eigenvalues of the matrix of second moments of area of the polygon around its principal axes [7]. The ESF for a circle is 1, and for an ellipse it is the ratio of major to minor axis length.

### 6.1. Node spacing ratio and volume change

In the continuous IBM, immersed boundaries are carried at precisely the local fluid velocity ((Equation 4)) because they are impermeable to fluid, in the sense that any given fluid particle will remain either inside or outside a particular immersed boundary for all time. In the discretized IBM, however, there is a gap of average length Δγ_*k*_ between any two adjacent nodes in boundary *k*. If this gap is much larger than the fluid mesh spacing, *h*, fluid flow between the nodes will have no impact on the propagation of node locations, and thus fluid will be able to flow across the boundary. Therefore, to ensure conservation of fluid volume within each immersed boundary, in the absence of fluid sources or sinks, Δγ_*k*_ must be small enough in relation to *h*, where the trade-off of making Δγ_*k*_ too small is simply computational expense.

To determine how small is small enough, Figure 4a shows the results of a set of simulations relating the change in volume of a circular immersed boundary to the node spacing ratio, Δγ_*k*_ /*h*. In each simulation, a circular cell is simulated for a fixed number of time steps. The intracellular spring properties are set with Δγ_*k*_ < *l*_*int*_ to ensure the linear springs are under tension and will, in the absence of the volume conservation property of the IBM, contract to reduce the perimeter of the immersed boundary. For each simulation we measure the proportional area change of the cell (the absolute change in area of the polygon divided by the original area), for a particular initial value of the node spacing ratio. From Figure 4a, we see that a node spacing ratio much above 2.0 results in poor volume conservation. A node spacing ratio of around 1.0, though, ensures that the numerical scheme matches the continuum limit well, while not being so small as to cause unnecessary computational overheads.

**Fig. 4.**
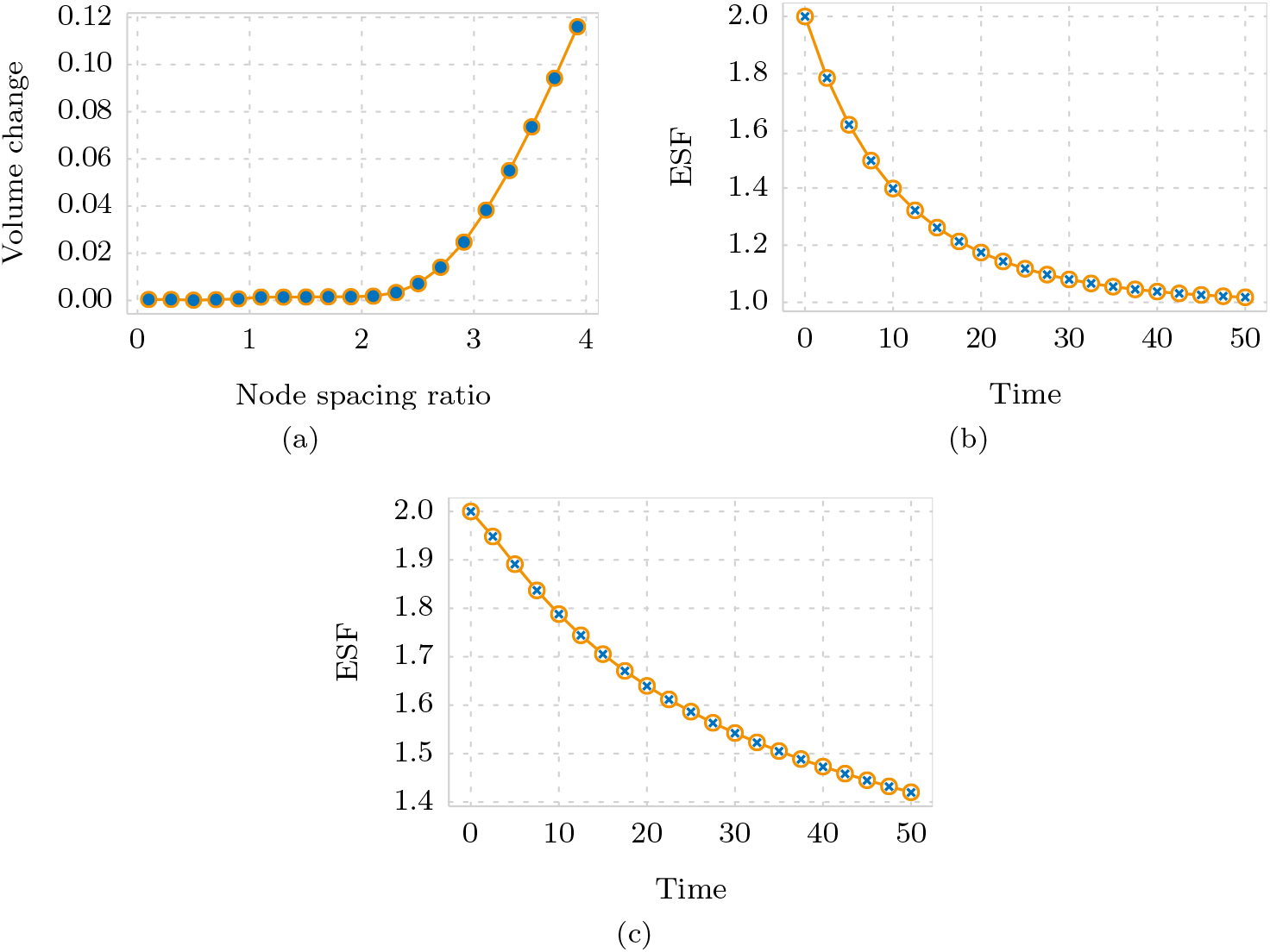
**(a) Node spacing ratio and volume change.** A set of simulations of a single circular immersed boundary, each run for the same fixed simulation time. Across the set of simulations the node spacing ratio, 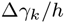, is varied and the proportional volume change of the immersed boundary is recorded. As the node spacing ratio increases beyond 2.0 there is a sharp increase in the proportional volume change, as a result of fluid escaping between the distantly spaced nodes. **(b) Scaling intra-cellular spring properties with node spacing.** Two simulations, each of an ellipse relaxing towards a circle, are run with the ESF sampled at 21 time points. Circles represent a simulation in which the immersed boundary is represented by N = 256 nodes, with intra-cellular spring constant 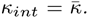. Crosses, coinciding with the circles, represent a simulation with a modified representation of N = 512 nodes and intra-cellular spring constant 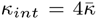. **(c) Scaling inter-cellular spring properties with node spacing.** Two simulations, each of two neighbouring ellipses relaxing, are run in which the ESF of one ellipse sampled at 21 time points. Circles represent a simulation in which the immersed boundary is represented by N = 256 nodes, with inter-cellular spring constant 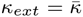. Crosses, coinciding with the circles, represent a simulation with a modified representation of N = 512 nodes and inter-cellular spring constant 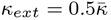, and with the intra-cellular spring properties scaled as in (b). For details on how to obtain the code for these simulations, which contains full details of all parameter values used, see Appendix A.

### 6.2. Scaling of individual cell properties

A single cell represented by an immersed boundary that is displaced out of equilibrium by, say, stretching, will relax back to a circle. If we were to run an identical simulation with half the time step, we would expect the dynamics to remain unchanged (up to numerical imprecision introduced as a result of the numerical scheme). Likewise, halving the uid mesh spacing, *h*, would, provided we obey the criteria of Subsection 6.1, leave the simulation output unchanged. Changing the immersed boundary representation, however, by altering the number of nodes per boundary, *N*_*k*_, requires a scaling of various parameters if we wish to recapitulate the same simulation.

To investigate this interplay, we consider the case where the node spacing in a single immersed boundary is decreased by a factor of α, starting from a reference value. Our goal is to derive the scaling required to ensure that the fluid flow, which determines the dynamics, remains unchanged. Two effects come in to play. First, the node spacing, Δγ_*k*_, which appears explicitly in the discretized force relation Equation (21), is reduced by a factor α, and therefore **F** must be increased by this factor in order to compensate. Second, since the boundary is represented by linear springs, we are now considering a system with *α* times the number of springs, each with length reduced by a factor *α*. Assuming the rest length, *l_int_*, scales proportionally with the length of the connection, the average energy of a spring in the reference configuration is given by

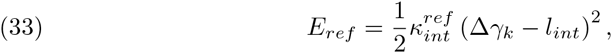

whereas the average energy of a spring in the new configuration is given by

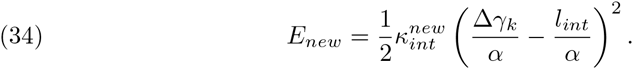

To ensure the potential in the immersed boundary is identical in both the reference and the new configurations, we must equate *E*_*ref*_ with α*E*_*new*_, giving 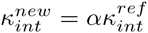. Combining the scaling by *α* from both considerations, we thus find that to increase the number of nodes in an immersed boundary by a factor α, we require an *α*^2^ increase in *k_int_*. Figure 4b verifies this scaling.

We now consider the case of two interacting cells with identical mechanical properties. If we alter the resolution of nodes around each immersed boundary, how must we change the cell-cell interaction force parameters *k*_*ext*_ and *l*_*ext*_ to recapitulate the same dynamics in a given simulation? Increasing the number of nodes by a factor α in each immersed boundary relative to a reference scenario will also increase the number of connections, determined via Equation (12), by a factor α. As the immersed boundaries are unchanged in size, *l*_*ext*_ should remain the same, and thus the potential contained within the boundary interactions will have increased in proportion to the number of connections. Thus, 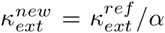 is the necessary scaling to ensure the simulation dynamics remain unchanged. Figure 4c shows summary statistics from a simulation verifying this scaling.

Putting these two results together, when increasing the density of nodes in a simulation by a factor *α*, we must scale 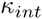 by *α*^2^, and 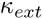 by 1/*α*. To encapsulate this within our computational framework, we introduce an `intrinsic length’ relative to which the scaling described here is applied. Due to this, the required scaling is not manually applied by the user; the simulation dynamics remain unchanged when the user alters the node spacing.

### 6.3. Convergence analysis

Here, we demonstrate how the numerical implementation converges with time step, fluid mesh spacing, and immersed boundary node spacing. We conduct this convergence analysis using a simple prototype simulation of an elliptical immersed boundary undergoing relaxation for a fixed simulation time. For each of the three parameters of interest, Δ*t*, *h*, and Δγ_*k*_, we perform a series of simulations where only the parameter of interest is varied, and collect a single summary statistic, the ESF, from which we can verify convergence.

To analyse convergence with time step, we run the relaxation simulation nineteen times, starting with Δ*t* = 0.5 and each time reducing Δ*t* by a factor of 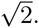 Figure 5 a demonstrates convergence of the ESF with time step. We assume the ESF associated with the finest time step to be the best approximation to the continuum limit, and define the error in ESF for each simulation to be the absolute difference between the ESF and this best value. Omitting the penultimate value, the gradient of a log-log plot of this error against time step is 1.11, demonstrating the order of convergence is approximately linear.

**Fig. 5.**
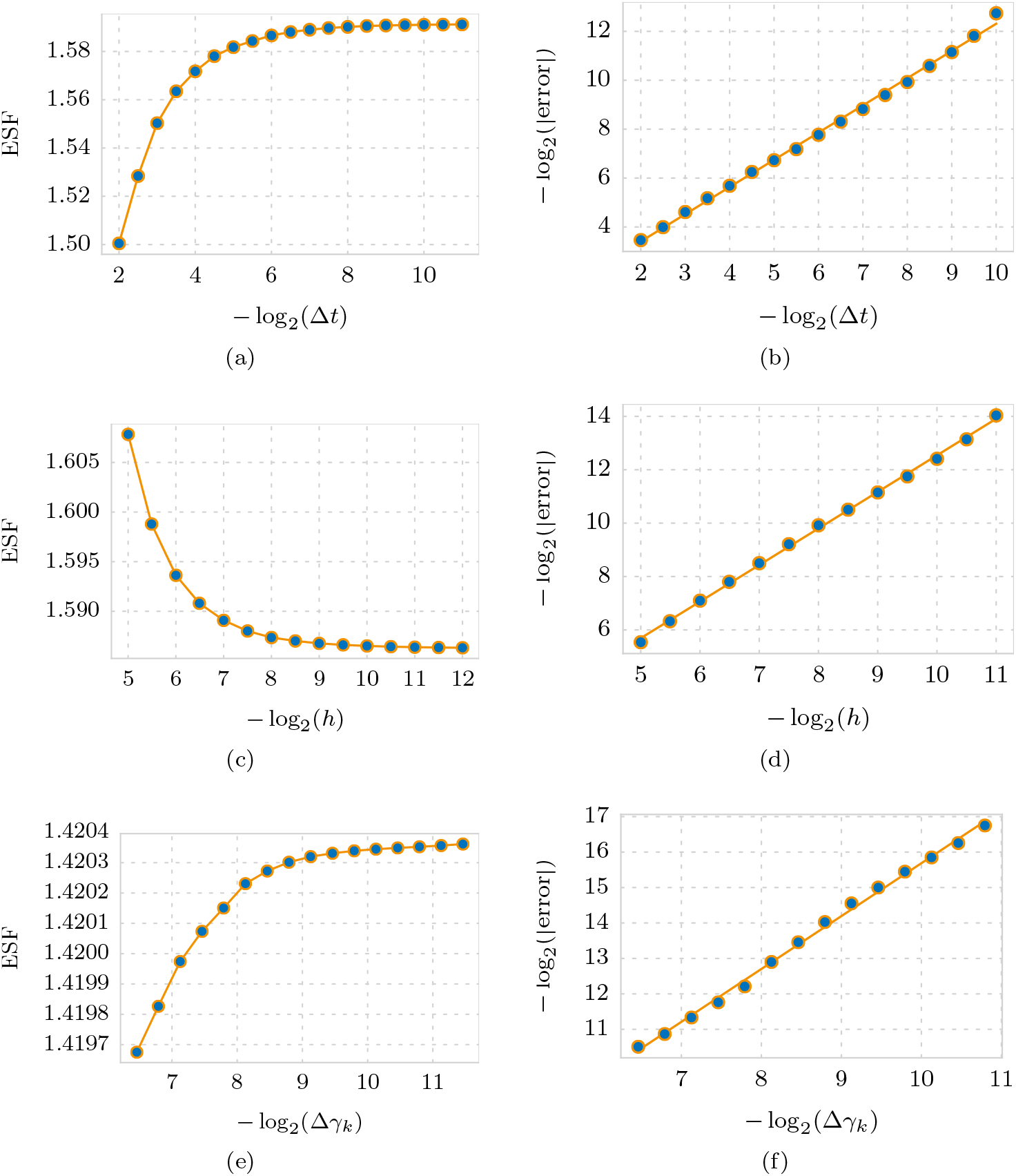
Convergence of computational implementation. (a) Convergence with time step. 19 simulations with different values of Δt were run for a fixed duration of 10 time units, with the following fixed parameters: initial ESF = 2.0, N = 128 nodes, l_int_ = 50% of node spacing, 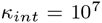, Re = 10^−4^, with 128×128 fluid mesh points, relative to an intrinsic spacing of 0.01. (b) Linear fit between error and time step, with a gradient of 1.11. (c) Convergence with fluid mesh spacing. 15 simulations with different fluid mesh spacings, h, were run, for a fixed duration of 10 time units, with the following fixed parameters: initial ESF = 2.0, N = 8192 nodes, l_int_ = 50% of initial node spacing, 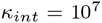, Re = 10^−4^, and Δt = 0.01, relative to an intrinsic spacing of 0.01. (d) Linear fit between error and fluid mesh spacing, with a gradient of 1.37. (e) Convergence with immersed boundary node spacing. 16 simulations with different numbers of immersed boundary nodes, therefore modulating Δγ_k_, were run for a fixed duration of 10 time units, with the following fixed parameters: initial ESF = 2.0, l_int_ = 50% of initial node spacing, 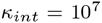, Re = 10^−4^, Δt = 0.01, and 64×64 fluid mesh points, relative to an intrinsic spacing of 0.01. (f) Linear fit between error and node spacing, with a gradient of 1.49. For details on how to obtain the code for these simulations, see Appendix A.

Similarly, to demonstrate convergence with fluid mesh spacing we run fifteen relaxation simulations starting with *h* = 1/32 and each time reducing h by a factor of 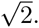. We need to pick a fixed large number of immersed boundary nodes to eliminate the node spacing ratio issue discussed in Subsection 6.2, and so as not to varyΔγ_*k*_. Figure 5c shows convergence of the ESF with *h*. Defining the error in a similar manner to above, we find the log-log gradient to be 1.37, demonstrating the order of convergence to be subquadratic. Finally, to demonstrate convergence with immersed boundary node spacing, we run sixteen relaxation simulations, starting with 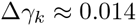 and each time reducing Δγ_*k*_ by a factor of 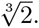. Figure 5 shows the ESF converging. The log-log gradient is 1.49, demonstrating the order of convergence to be subquadratic.

In addition to convergence of the numerical implementation, we also require our implementation of cell division to converge with immersed boundary node spacing: for a given cell division, the shape of the resulting daughter cells should be independent of the choice of boundary parametrisation. We verify this convergence by performing cell division operations on a number of elliptical immersed boundaries, each represented by a different number of nodes, and using the ESF as a summary statistic of daughter cell shape. Figure 6 shows results with a log-log gradient of 1.96, demonstrating the order of convergence to be quadratic.

**Fig. 6.**
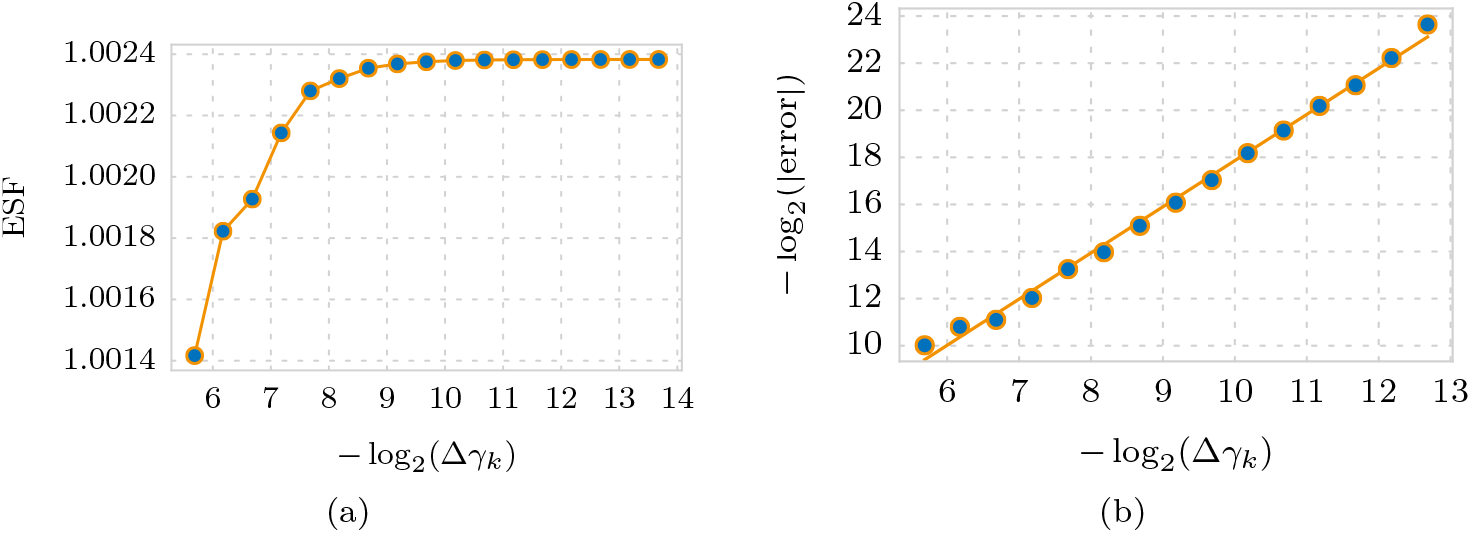
Convergence of cell division implementation. (a) 17 simulation results showing the ESF of an immersed boundary resuling from the application of the cell division algorithm (subsection 5.2), from elliptical immersed boundaries with varying values of Δγ_*k*_. (b) Linear fit between error and Δγ_*k*_, with a gradient of 1.96. For details on how to obtain the code for these simulations, see Appendix A.

## 7. Discussion

In this manuscript, we have presented a thorough description of the equations governing the IBM, and full details of a common discretisation approach and method of numerical solution. We have presented an efficient computational implementation, as part of a mature and thoroughly tested C++ library designed specifically for computational biology simulations. We have demonstrated numerically various parameter scaling properties of the IBM, and have demonstrated the convergence properties of our implementation. In this section, we return to several choices made during the formulation of our IBM model.

### 7.1. Stokes or Navier-Stokes

The first such choice was whether to solve the full Navier-Stokes equations, or whether to solve the Stokes equations in the low Reynolds Number limit. To address this question, we first emphasize that the `fluid’ underlying the IBM need not have a direct physical correlate. It may be helpful to think of the fluid simply as a tool by which the positions of the boundaries are updated, and which has certain `nice’ properties (such as volume preservation inside closed contours), although some authors have nevertheless sought to draw parallels between this fluid and the cell cytoplasm and extracellular medium [23]. A concrete example, though, of the difference between the IBM fluid and the fluid-like properties of the underlying biological system is in the case of a stationary circular boundary: if there is a resultant elastic force, there will be a non-zero body force in the IBM fluid and therefore an induced flow. Since the fluid cannot be assumed to faithfully represent underlying biology, it is not obvious that modelling a biological situation with small Reynolds number necessarily means the Reynolds number in an associated immersed boundary problem need also be small. Rejniak et al., for instance, derive a `biological’ Reynolds number of 10^−9^, but use the value 5.9×10^−5^ for their simulations [24]; a value chosen so as to recapitulate the relevant dynamics. This discrepancy demonstrates that the IBM fluid cannot be expected to adequately mimic the fluid-like properties of the underlying biology, and thus that we must take care in assuming an appropriate Reynolds number in IBM simulations need necessarily be very small. Further investigation is required to ascertain the relationship between `fluid’ properties in vivo and in silico for the IBM. Cutting experiments, for instance, where tissue is observed to recoil after ablation, could be used to fit an appropriate Reynolds number for the IBM in order to match in vivo dynamics. While IBM implementations based on Stokes flow do exist [2, 12, 30], we have chosen to implement the full generality of the Navier-Stokes problem. This keeps open the possibility of modelling situations where inertial effects cannot necessarily be neglected, while acknowledging that there are scenarios in which the reduced problem may be appropriate, and computationally much less expensive to solve.

### 7.2. Discrete delta function

We made a specific choice for the form of the discrete delta function. Peskin [17] derived the following form for *ϕ*, in contrast to that presented in Equation (10):

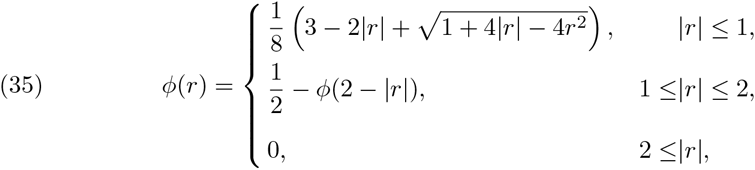

While the functional form appears quite different, the numerical values taken by the different formulations of *ϕ* are very similar (differing by less than 0.008 at any point in the domain). Given this incredibly similarity, using one form rather than the other may be decided by computational efficiency. In practice, we find the trigonometric function slightly quicker to compute during a simulation, which is likely due to difficult branch prediction of the `if’ statement necessary to compute *ϕ* using Equation (35). The proportion of the total simulation time spent evaluating the discrete delta function is, however, small enough that in practical terms the choice of *ϕ* is immaterial.

### 7.3. Inter-cellular interaction terms

Third, we will briefly discuss the choice of functional form for the inter-cellular interaction terms. The sharp cut-off represented by the interaction distance *d*_*ext*_ in Equation (12) may be unphysical, as it implies that when boundaries move apart, the opposing force linearly increases with distance until instantaneously becoming zero at distance *d*_*ext*_. A different functional form may mirror the underlying behaviour more closely, and one such example is the Morse potential [15], which has a functional form

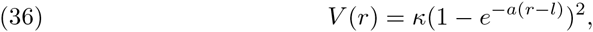

where *κ* and *a* denote the depth and width of the potential well, respectively, *r* is the distance between the interacting nodes, and *l* is the equilibrium distance of the bond. The force between two immersed boundary nodes would, as a result of such a potential, be exponentially repulsive at short distances, have an attractive peak at a medium distance, and tail off at long distance. A cut-off at a value of *d_ext_* would still be needed for computational reasons, but this cut-off would be at a low value of the force rather than at the maximum value, as is the case with linear springs. To what extent the choice of functional form impacts immersed boundary simulations is an open question, and a topic for further study.

### 7.4. Balancing sources

Finally, in Subsection 4.7 we gave no precise formulation for the number of fluid sources, *M* − *N*, in excess of those associated to immersed boundaries. The purpose of these additional sources is to balance the net fluid creation due to processes such as cell growth, to ensure a constant fluid volume within the domain Ω. In our implementation we choose *M* ≈ 2*N*, and initially place these equidistant along the boundary *y* = 0. Rejniak and colleagues [24] use a similar approach, but do not specify the number of such additional sources, while Dillon and Othmer [5] use exactly four, but do not specify their initial locations. The implications of such choices have not been systematically investigated, and to what extent these choices impact upon the results of simulations is a topic for further study.

## 8. Conclusion

Through the availability of ever richer datasets from molecular and live-imaging studies, we are in a position to undertake data-driven computational modelling of morphogenetic processes. In tandem, the ready availability of computing power allows not only for individually costly simulations to be run, but also for parameter estimation or sensitivity analysis studies requiring thousands of such simulations. The time is ripe, therefore, to take advantage of both the accessibility of high-resolution data and the availability of enormous computational power. We have presented here an open-source, efficient, and modular implementation of the IBM, one such framework able to make use of both.

A strength of such models is the ease with which cellular heterogeneity (for example, through patterning mechanisms) may be incorporated, and the consequences for tissue-scale behaviour be simulated and explored. The development of methods to efficiently explore the parameter space of such models, perform inference and model calibration against quantitative datasets, and analyse the tissue-level mechanical properties of such models remain avenues for future work in this area.

## Appendix A. Obtaining the source code

The C++ implementation of the IBM within Chaste is available as a feature branch as part of the publically accessible Chaste Git repository. Details on accessing this repository can be found at https://chaste.cs.ox.ac.uk/trac/wiki/ChasteGuides/GitGuide, and the IBM branch is titled `fcooper/immersed boundary’.

The code used for simulations in this paper is provided as a zipped folder (see Supplementary Material). This code takes the form of a `user project’ entitled `Ib-NumericsPaper’ which can be interfaced with Chaste using instructions at https://chaste.cs.ox.ac.uk/trac/wiki/InstallGuides/CheckoutUserProject.

Within this user project, numerical convergence simulations can be found in /apps/src, where ‘.cpp’ files define the simulations and ‘.py’ _les run the simulations and perform the post-processing. Profiling simulations are defined in the test suite /test/TestPro_ling.hpp. All other simulations are defined as individual tests, which are defined and documented in the test suite /test/TestNumericsPaperSimulations.hpp.

## Acknowledgements

The Chaste developers have been of great and varied help. In particular, we wish to thank (in alphabetical order) Jonathan Cooper, Jochen Kursawe, Gary Mirams, Joe Pitt-Francis, and Martin Robinson.

## Footnotes

1 http://www.cs.ox.ac.uk/chaste/download.html

2 http://cxxtest.com/

## Notes

* The first author’s work was supported by funding from the Engineering and Physical Sciences Research Council (EPSRC) [grant number EP/G03706X/1].

